# Organic Fertilizer Substitution for Chemical Fertilizer Improves Yield, Medicinal Active Compounds and Amino-Acid Profiles of *Dendrobium officinale* with Multi-Index Comprehensive Evaluation

**DOI:** 10.64898/2026.10.29.762708

**Authors:** Long Zhou, Wenna Zhou, Xudong Zhao, Huarong Liu, Hongbin Yang

## Abstract

*Dendrobium officinale* (*D. officinale*) is an important medicinal horticultural crop. Its wild resources are threatened by over-exploitation. Fertilization regimes that balance yield and medicinal quality are urgently required for its artificial cultivation. This study aimed to investigate the effects of reduced chemical fertilizer combined with organic fertilizer on the yield and quality of *D. officinale*, and screen the optimal combined fertilization pattern, so as to provide a theoretical basis for standardized fertilization in *D. officinale* cultivation. A bed-cultivation experiment was carried out in greenhouse. Seven treatments were set up, including no fertilizer application (CK), sole chemical fertilizer (CF), 20% organic fertilizer substitution for chemical fertilizer (T20), 40% organic fertilizer substitution for chemical fertilizer (T40), 60% organic fertilizer substitution for chemical fertilizer (T60), 80% organic fertilizer substitution for chemical fertilizer (T80), and single organic fertilizer application (OM). Plant growth traits, yield, active ingredients, protein and amino acid components of *D. officinale* were determined. The membership function method was adopted for comprehensive evaluation of all treatments. Fertilization significantly improved the growth traits, yield and quality of *D. officinale*. Compared with CK, stem length, stem diameter and yield under fertilized treatments increased significantly by 15.5%-60.3%, 34.2%-64.7% and 65%-143.2%, respectively; dendrobine, polyphenols and alcohol-soluble extract contents increased significantly by 21.2%-83.7%, 14.0%-49.1% and 10.8%-26.9%, respectively. The highest plant growth and yield were observed under sole chemical fertilizer treatment. With the increase of organic fertilizer substitution ratio, plant growth and yield decreased gradually, while medicinal active ingredients including dendrobine, polysaccharides and flavonoids showed a trend of rising first and then falling. A total of 17 amino acids were detected in all treatments. T40 and T60 treatments significantly increased total amino acids, essential amino acids, as well as contents of medicinal, umami and bitter-tasting amino acids, whereas excessive proportion of organic fertilizer led to the decline of amino acid contents. Most growth, quality and amino acid indicators showed significant or extremely significant positive correlations with each other. The comprehensive evaluation order by membership function method was T40 > T60 > CF > T20 > T80 > OM > CK, and T40 obtained the highest comprehensive evaluation value. Sole chemical fertilizer contributes to yield increase of *D. officinale*, while combined application with organic fertilizer improves its medicinal quality. The 40 % organic substitution scheme provides a feasible fertilization strategy for high-quality greenhouse production of this medicinal horticultural plant.

## 1. Introduction

*D. officinale*, a perennial epiphytic herb belonging to Orchidaceae Dendrobium genus, is a precious medicinal horticultural crop in China. It has attracted extensive attention in pharmaceutical and agricultural research owing to its unique medicinal efficacy and abundant nutritional components[1]. Its medicinal value has been well-documented in ancient Chinese herbal classics. Compendium of Materia Medica records that it could “replenish yin and essence, and tonify the stomach and intestines upon long-term consumption”. Ranked first among the legendary “Nine Immortal Herbs” of traditional Chinese medicine and reputed as “life-saving immortal herb” among folk communities, *D. officinale* is rich in bioactive substances including polysaccharides, alkaloids, polyphenols and flavonoids. Modern pharmacological studies have verified its multiple pharmacological activities such as antioxidant capacity, immunomodulation, anti-tumor effect and blood-pressure-lowering and vascular-protective functions, endowing it with high medicinal and health-care values[2].

*D. officinale* possesses complex nutritional profiles and outstanding nutritional value, and abundant mineral elements as well as functional nutrients constitute the material basis for its health-promoting effects[3]. Dendrobium polysaccharides, alcohol-soluble extracts and dendrobine are core quality indicators for evaluating medicinal materials of *D. officinale* and play vital roles in medical and health-care applications[4]. As the primary functional component, dendrobium polysaccharides can effectively enhance organismal immunity, regulate blood glucose, exert anti-oxidation and anti-aging effects, and inhibit the proliferation of tumor cells[5]. Alcohol-soluble extracts mainly consist of polyphenols, flavonoids, alkaloids and other active compounds. Among them, polyphenols and flavonoids exhibit remarkable free-radical scavenging capacity to delay cellular senescence, together with physiological functions including anti-inflammation, anti-biosis and immunoregulation[6]. Dendrobine, a characteristic alkaloid specific to *D. officinale*, exerts prominent effects in anti-fatigue, memory improvement, organism development promotion and anti-tumor activities[7]. Furthermore, *D. officinale* contains multiple essential amino acids that human bodies cannot synthesize independently. Components such as glutamic acid, aspartic acid, leucine and alanine serve as critical material carriers for its clinical health-care efficacy. They participate in diverse human physiological metabolic processes and perform irreplaceable functions in maintaining physical health, facilitating growth and development, and strengthening immune functions[8−10]. Accordingly, systematic investigation into the composition and concentration characteristics of nutritional and active components in *D. officinale* can not only clarify its medicinal and health-care mechanisms, but also provide theoretical support for the scientific exploitation and efficient utilization of *D. officinale* resources[11].

Benefiting from its superior medicinal and economic values, the market demand for *D. officinale* keeps increasing. Nevertheless, long-term excessive wild collection, destruction of natural habitats and low reproduction coefficient of wild populations have led to sharp decline of its wild natural resources. Wild *D. officinale* is now listed as critically endangered and can hardly satisfy the requirements of industrial development[12]. Artificial cultivation acts as a core approach for germplasm conservation of wild *D. officinale* and resolution of market supply-demand contradiction, and it represents the dominant pattern for current industrial production of Dendrobium. With continuous innovation in cultivation techniques, greenhouse seedling raising and large-scale planting have been widely popularized, effectively solving industrial bottlenecks such as seedling shortage and low transplant survival rate, and laying foundations for large-scale production of *D. officinale*[13]. However, practical production has revealed a prevalent problem in artificially-cultivated *D. officinale*: yield increases while quality degrades. The content of bioactive components in cultivated medicinal materials is markedly lower than that in wild counterparts, which seriously impairs its commodity value and restricts the high-quality development of *D. officinale* industry. Hence, exploring scientific cultivation regulation technologies to realize synergistic improvement in yield and quality of artificially-cultivated *D. officinale* has become a key and challenging issue in current Dendrobium cultivation research.

Diverse artificial cultivation patterns have been developed for *D. officinale*, mainly including greenhouse bed cultivation, under-forest epiphytic cultivation on living trees, and simulated-wild epiphytic cultivation on rocks. Different cultivation modes, environmental conditions and management practices significantly affect growth-metabolism, accumulation of bioactive ingredients and medicinal quality of *D. officinale*[14-18]. Existing studies have demonstrated that *D. officinale* under under-forest tree-epiphytic cultivation presents significantly higher contents of polyphenols, flavonoids, free amino acids and in-vitro antioxidant activities than those under greenhouse pot cultivation and board-attached cultivation[19]. Higher accumulation of flavonoids and flavonols is observed under rock-epiphytic and tree-epiphytic systems compared with greenhouse cultivation, whereas rock-epiphytic samples show advantages in polysaccharide content and antioxidant activity[20]. Meanwhile, researchers have carried out regulatory experiments on environmental factors including cultivation substrate, planting cycle and light intensity, confirming that appropriate cultivation conditions can effectively boost plant growth and development and optimize quality indicators of *D. officinale*.

Apart from environmental factors, exogenous substance regulation and fertilization management are important strategies to promote plant growth and improve quality of *D. officinale*. Liu et al.[21] reported that exogenous application of 2,3-butanediol fermentation broth, γ-aminobutyric acid, melatonin and other substances could simultaneously promote biomass accumulation as well as the synthesis of polysaccharides and alkaloids in protocorms of *D. officinale*. In terms of fertilization regulation, available researches mainly focus on single fertilization and foliar fertilization modes. Spraying nano-calcium fertilizer facilitates the accumulation of endogenous phytohormones and soluble sugars, and accelerates stem elongation and tillering of *D. officinale*[22]. Application of botanical fertilizer markedly enhances antioxidant capacity and promotes polysaccharide accumulation[23]. Special foliar fertilizers and combined organic-inorganic foliar fertilizers can improve growth parameters and quality traits of *D. officinale* to varying degrees, and optimal fertilizer concentration maximizes fertilization benefits. Fertilization has been proven as a critical measure regulating yield and quality of *D. officinale*. However, current researches mostly concentrate on single fertilizer application and foliar fertilization. Systematic studies on chemical fertilizer reduction and scientific combined application of organic and chemical fertilizers remain scarce. Particularly, fertilization ratio schemes balancing high yield and superior quality of *D. officinale* are insufficient, which cannot meet the production demands for standardized and high-quality cultivation[24].

Current studies concerning quality improvement of *D. officinale* predominantly focus on environmental regulation such as cultivation pattern, substrate, light and growth cycle, and relatively sound technical systems for cultivation optimization have been established[25-26]. Nevertheless, research gaps still exist in fertilization regulation for synergistic promotion of yield and quality.

Fertilization constitutes a core field management measure for yield increase and quality optimization of crops. Combined application of organic and inorganic fertilizers is an efficient fertilization mode to stabilize crop yield and optimize quality, which has been validated in multiple cash crops. However, its application in greenhouse-cultivated *D. officinale* has rarely been reported. Few studies have systematically evaluated how organic substitution gradients alter not only yield and secondary metabolites, but also comprehensive amino-acid profiles (including medicinal- and umami-taste amino acids) for greenhouse-bed cultivated *D. officinale*. Multi-indicator comprehensive assessment balancing agronomic yield and nutraceutical quality remains insufficient.

Greenhouse bed-cultivated *D. officinale* was adopted as experimental material in this study. Treatments with different gradients of organic manure substituting chemical fertilizer were set up to systematically investigate the effects of reduced chemical fertilizer combined with organic manure on growth-yield, medicinal bioactive components and amino-acid profiles of *D. officinale*. The optimal fertilization ratio was screened out through multi-indicator comprehensive evaluation. This study aims to clarify the regulatory effects of combined organic-chemical fertilization on yield and quality of *D. officinale*, identify fertilization patterns balancing high yield and good quality, and provide scientific fertilization schemes as well as theoretical foundations for standardized high-quality greenhouse cultivation of *D. officinale*, so as to contribute to quality-benefit improvement and green sustainable development of Dendrobium industry.

## 2. Materials and Methods

### 2.1 Overview of the Experimental Site

The experiment was carried out in a standardized multi-span greenhouse at the National Dendrobium Germplasm Resource Bank in Ruili, Dehong Prefecture, Yunnan Province (23°52′26″ N, 97°56′31″ E). Located in the tropical monsoon climate zone of western Yunnan, the site sits at an altitude of 780 m, with an annual average temperature of 21.3 °C and annual precipitation of 1450 mm. It is frost-free year-round, with sufficient solar radiation and favorable humidity, representing a genuine producing area suitable for *D. officinale* cultivation.

The greenhouse features controllable environmental conditions: annual indoor temperature ranges from 18 °C to 28 °C, relative air humidity is maintained at 75%-90%, and the shading rate is set at 70%. With good ventilation, free from industrial pollution and low risk of severe pest and disease outbreaks, the greenhouse meets the standards for germplasm conservation and standardized cultivation trials of Chinese medicinal materials.

The cultivation substrate consisted of decomposed pine bark, coconut husk chips and volcanic rock at a volume ratio of 6:3:1 (v/v). The substrate was fully decomposed, free of insect eggs and mildew, and disinfected by sunlight exposure prior to use. Its basic physicochemical properties were as follows: pH 5.5-6.2, organic matter 42.6 g·kg^−1^, total nitrogen 1.83 g·kg^−1^, available phosphorus 36.5 mg·kg^−1^, and available potassium 128.4 mg·kg^−1^. Characterized by loose texture, good aeration and favorable water-fertilizer retention capacity, the substrate received no fertilizer application before transplantation and exhibited uniform fertility, so as to eliminate the interference of background fertility differences across experimental units.

### 2.2 Experimental Materials

1-year-old acclimated tissue-cultured plantlets of *D. officinale* were obtained from the National Dendrobium Germplasm Resource Bank in Ruili. The plantlets had pure genetic background, uniform growth performance, and were free of pests, diseases and mechanical damage. Each seedling was 8-10 cm high with 2-3 valid tillers. Uniform-sized plantlets were selected for transplantation to guarantee consistency of experimental materials.

The tested organic fertilizer was a special organic fertilizer independently developed by the base (CN202510102937.8). It was fermented with wormcast, *Pleurotus eryngii* mushroom residue and mugwort straw as raw materials. The fermentation substrate was soaked in aqueous solution containing *D. officinale* extract and mugwort extract, followed by inoculation with *Bacillus subtilis* strain H728 for thorough decomposition. This organic fertilizer contained 2.5% total nitrogen with a moisture content of 28.0%. The chemical fertilizer adopted was a balanced compound fertilizer (N-P_2_O_5_-K_2_O 15-15-15, total nutrient ≥45%), purchased from Yunnan Yuntianhua Co., Ltd.

### 2.3 Experimental Design

*D. officinale* was cultivated on elevated growing beds approximately 60 cm above the ground inside the greenhouse. Each bed measured 12 m × 1.5 m, with a 60 cm spacing between adjacent beds. A single-factor randomized block design was adopted in this experiment, with equivalent nitrogen supply as the core principle for organic fertilizer substitution. Seven treatments of combined organic-inorganic fertilization were set: no fertilizer application (CK), sole chemical fertilizer (CF), 20% organic fertilizer substitution for chemical fertilizer (T20), 40% organic fertilizer substitution for chemical fertilizer (T40), 60% organic fertilizer substitution for chemical fertilizer (T60), 80% organic fertilizer substitution for chemical fertilizer (T80), and sole organic fertilizer (OM). Each treatment was replicated three times, generating a total of 21 experimental plots. Each plot covered an area of 4 m × 1.5 m = 6 m^2^, where 171 clumps of *D. officinale* were planted at a planting density of 15 cm × 20 cm with consistent planting depth.

The balanced compound fertilizer served as the reference fertilizer for equivalent-nitrogen treatment. The application rate of organic fertilizer was calculated based on its total nitrogen content and moisture content. The annual nitrogen application rate for *D. officinale* was set at 180 kg·ha^-1^. The full amount of organic fertilizer was thoroughly mixed with the cultivation substrate and applied as basal fertilizer by layered broadcasting; transplantation was performed after substrate maturation. Chemical nitrogen fertilizer was applied four times throughout the growing cycle, accounting for 40%, 15%, 30% and 15% of total nitrogen at basal stage, sprouting stage, vigorous-growth stage and quality-accumulation stage respectively, and uniform irrigation scheduling was applied for all experimental plots to maintain substrate moisture.

Uniform field management including watering, ventilation, shading, green pest-disease control and weeding was implemented for all treatments. No foliar fertilizer or plant growth regulators were sprayed during the whole experimental period. The trial lasted for one complete growth cycle (March to February of the subsequent year). Details of fertilization treatments are shown in Table 1.

**Table 1.** Experimental Treatments.

| Treatment | Fertilization mode | Fertilizer application amount(kg·ha<sup>-1</sup>) |  |
|---|---|---|---|
|  |  | Compound fertilizer(15-15-15) | Organic fertilizer(2.5%N) |
| CK | No fertilization | 0 | 0.0 |
| CF | Sole chemical fertilizer | 1200 | 0.0 |
| T20 | 20% organic fertilizer + 80% chemical fertilizer | 960 | 2000.0 |
| T40 | 40% organic fertilizer + 60% chemical fertilizer | 720 | 4000.0 |
| T60 | 60% organic fertilizer + 40% chemical fertilizer | 480 | 6000.0 |
| T80 | 80% organic fertilizer + 20% chemical fertilizer | 240 | 8000.0 |
| OM | Sole organic fertilizer | 0 | 10000.0 |

**Table 1.** Effects of different fertilization treatments on growth traits of *D. officinale*.

| Treatment | Stem length (cm) | Stem diameter (mm) | Internode length (cm) | Number of leaves (leaves Cluster<sup>-1</sup>) | Chlorophyll (mg kg<sup>-1</sup>) | Number of fresh shoots (Branch Cluster<sup>-1</sup>) | Yield (kg ha<sup>-1</sup>) | Dry weight ratio (%) |
|---|---|---|---|---|---|---|---|---|
| CK | 36.23±2.28e | 3.16±0.09f | 1.33±0.04e | 151.5±3.7e | 4.25±0.09e | 10.5±0.7e | 3487.5±130.1f | 23.82±0.39c |
| CF | 58.1±1.76a | 5.2±0.06a | 2.12±0.14a | 318.6±4.3a | 5.79±0.03a | 21.9±1.2a | 8482.5±170.2a | 25.53±0.28a |
| T20 | 54.57±0.9a | 4.6±0.04b | 1.92±0.05a | 294.7±6.6b | 5.34±0.11b | 19.5±0.7a | 8130±73.2b | 24.62±0.16b |
| T40 | 52.47±0.8b | 4.52±0.05bc | 1.8±0.03b | 275.4±6.6b | 5.14±0.1b | 17.7±0.5b | 7999.5±108.6b | 24.4±0.06bc |
| T60 | 49.43±1.06c | 4.43±0.06cd | 1.67±0.03c | 263.7±4.4c | 4.87±0.1c | 14.7±0.2c | 7427.5±127.8c | 24.72±0.25b |
| T80 | 46.2±0.6c | 4.39±0.04d | 1.54±0.05d | 241.9±7d | 4.56±0.11c | 14.4±0.6c | 6714.5±109.6d | 24.09±0.67bc |
| OM | 41.87±1.46d | 4.24±0.07e | 1.47±0.03d | 248.5±7.8d | 4.51±0.11d | 12.6±0.5d | 5759.5±148.8e | 24.52±0.38b |

Note: CK: No fertilization; CF: Sole chemical fertilizer; T20: 20% organic fertilizer + 80% chemical fertilizer; T40: 40% organic fertilizer + 60% chemical fertilizer; T60: 60% organic fertilizer + 40% chemical fertilizer; T80: 80% organic fertilizer + 20% chemical fertilizer; OM: Sole organic fertilizer. Values are means ± standard deviation (*n* = 3). Values with the same lower-case letters are not significantly different among different treatments at the 5% level by the LSD. The same below.

### 2.4 Determination of Measurement Indicators

#### 2.4.1 Determination of Growth Indices

Plant-growth traits and yield indices were measured after one complete annual planting cycle of *D. officinale*. The five-point sampling method was adopted. Five sampling points were randomly set within each experimental plot, and three clumps of healthy *D. officinale* plants with uniform growth and free of pests and diseases were selected at each point as test samples. Growth indices including stem length, stem diameter, internode length, number of current-year fresh stems, chlorophyll content and yield were measured and recorded.

Specific measurement procedures were as follows: Stem length per clump was measured with a meter ruler from the second internode at the base of pseudobulb to the apex of pseudobulb. Stem diameter was determined at the middle part of the fourth internode upward from the pseudobulb base using a vernier caliper to guarantee consistent measuring position and reliable data. The total number of internodes and leaves per clump were manually counted, and the average internode length was calculated by the formula:

Internode length = stem length / number of internodes.

The number of newly-emerged stems in the current year was counted and recorded as the number of fresh stems. Chlorophyll content of functional leaves was determined by a SPAD-502 portable chlorophyll meter, with three replicate measurements per plant for averaging.

After determination of morphological indices, the above-ground stems and leaves of *D. officinale* were harvested. Total fresh weight of samples was accurately weighed and recorded using an electronic balance. Fresh samples were divided into two subsamples for subsequent treatments. The first subsample was cut into pieces and mixed evenly, subjected to fixation at 90 °C for 15 min in an oven, and then dried to constant weight at a constant temperature of 60 °C. Dry-sample weight was recorded. Water content was calculated according to the formula:

Water content (%) = (fresh weight − dry weight) / fresh weight × 100%.

The dried samples were ground, sieved and stored in sealed containers for subsequent quality-index determination. The second set of fresh stem samples were packed into sampling bags, snap-frozen in liquid nitrogen for 30 min, and preserved in an ultra-low-temperature refrigerator for the assay of medicinal bioactive components.

### 2.4.2 Determination of Quality Indices

Quality indices were assayed using the two sets of preserved *D. officinale* samples mentioned above. Three technical replicates were arranged for each index. Detailed determination methods and calculation formulas are described below.

Alcohol-soluble extracts (AE) were determined in accordance with the standard method specified in Volume I of *Pharmacopoeia of the People’s Republic of China* (2020 Edition)[27].

Contents of polysaccharides (Po) and total flavonoids (TF) were measured by ultraviolet spectrophotometry following Wu et al.[28]. The unified calculation formula was:

Index content (mg·g^-1^) = C × V / M,

where C = concentration of sample solution derived from standard-curve fitting; V = total volume of sample extract; M = actual weighed mass of sample. Rutin was used as the reference substance for total-flavonoid determination.

Soluble-protein content was determined via Coomassie brilliant blue method described in Volume III of *Pharmacopoeia of the People’s Republic of China* (2020 Edition)[29].

Total alkaloids (AL, expressed as dendrobine equivalent) were assayed by acid-dye colorimetry according to Ao et al.[30]. The formula was:

Alkaloid content (mg·g^-1^) = C × (V ÷ V_1_) ÷ W,

where C = dendrobine concentration in reaction solution (μg); V = total volume of sample extract (mL); V_1_ = volume of reaction solution for measurement (mL); W = mass of tested sample (g).

Crude fiber (CF) was measured following the national standard GB/T 5009.10-2003[31]. The formula was: Crude-fiber content (%) = M / m × 100%,

where m = mass of tested sample (g); M = mass of residue or mass loss after high-temperature ashing (g).

Total polyphenol content was determined by ultraviolet spectrophotometry with gallic acid as reference substance, as described by Huang et al.[6], with minor optimization of sample pretreatment. The calculation formula was consistent with that for total flavonoids:

Polyphenol content (mg/g) = C × V / M, and parameters were defined as above.

A total of 17 amino-acid species were quantified referring to the national food-safety standard GB 5009.124-2016 *National Food Safety Standard-Determination of Amino Acids in Foods*[32]. According to biosynthetic origin and human nutritional requirements, the 17 amino acids were classified into essential amino acids (Threonine-Thr, Phenylalanine-Phe, Lysine-Lys, Valine-Val, Methionine-Met, Isoleucine-Ile, Leucine-Leu) and non-essential amino acids (Aspartic acid-Asp, Tyrosine-Tyr, Serine-Ser, Glutamic acid-Glu, Glycine-Gly, Alanine-Ala, Cystine-Cys, Histidine-His, Arginine-Arg, Proline). Based on physiological properties and functions, amino acids were further divided into umami-taste amino acids (4 species: Asp, Glu, Gly, Ala), bitter-taste amino acids (8 species: Val, Met, Ile, Leu, Tyr, Phe, His, Arg) and medicinal amino acids (9 species: Asp, Glu, Gly, Arg, Met, Phe, Leu, Lys, Tyr). The nutritional differences in amino-acid profiles among fertilization treatments were reflected by calculating their respective mass fractions. The formulas were as follows:

Essential-amino-acid fraction (EAA/TAA, %) = content of essential amino acids / total amino-acid content Non-essential-amino-acid fraction (NEAA/TAA, %) = content of non-essential amino acids / total amino-acid content Medicinal-amino-acid fraction (MAA/TAA) = content of medicinal amino acids / total amino-acid content

### 2.5 Comprehensive Evaluation of Growth and Quality of D. officinale

The membership-function method[33-34] was applied for comprehensive evaluation of growth and quality of *D. officinale* under different fertilization treatments.

For positively-correlated indices, membership-function values {*u*(*X*_*j*_)} were calculated using Equation (1).

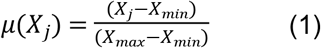

For indices negatively correlated with plant growth and quality, membership-function values were calculated using Equation (2).

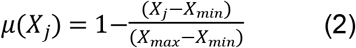

Standard coefficients(*V*_*j*_) and weights(*W*_*j*_) for each index were computed via Equation (3) and Equation (4).

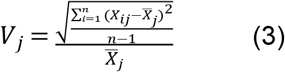

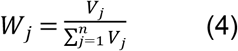

The comprehensive evaluation value (*D*) was obtained by Equation (5).

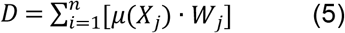

Where:

*X*_*j*_ represents the measured value of the j-th index; *X*_*min*_ represents the minimum value of the j-th index; *X*_*max*_ represents the maximum value of the j-th index;

*X*_*ij*_ represents the membership-function value of the j-th index under the i-th treatment;

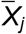 represents the average membership-function value of the j-th index across all treatments;

*D* represents the comprehensive evaluation value; higher D values indicate better growth performance and higher medicinal quality of *D. officinale*;

*n* represents the renumber of experimental treatments.

### 2.6 Statistical Analysis and Graphing

Raw experimental data were preliminarily processed in Excel 2020. One-way analysis of variance (ANOVA) and correlation-analysis computation were performed using IBM SPSS Statistics 27. Duncan’s multiple-range test was adopted for post-hoc comparison. Origin 2021 was used for figure plotting.

## 3. Results

### 3.1. Effects of Different Fertilization Treatments on Growth Traits of *D. officinale*

Different fertilization treatments exerted significant effects on the growth traits of *D. officinale* (Table 1). Compared with the non-fertilization control (CK), sole chemical fertilizer application, reduced chemical fertilizer combined with organic manure, and sole organic fertilizer application all significantly modified stem length, stem diameter, internode length, leaf number, chlorophyll content, number of fresh stems and yield of *D. officinale*. These parameters increased by 15.5%-60.3%, 34.2%-64.7%, 10.3%-59.0%, 59.7%-110.3%, 6.0%-36.1%, 20.1%-108.5% and 65.0%-143.2%, respectively, relative to the non-fertilization treatment (*P*<0.05). Compared with CK, the dry-matter conversion rate of CF, T20, T60 and OM treatments increased significantly by 7.2%, 3.4%, 3.8% and 3.0%, respectively (*P*<0.05).

Among all fertilization regimes, the sole chemical-fertilizer treatment (CF) achieved the optimal performance in growth traits and yield of *D. officinale*. Treatments with chemical-fertilizer reduction combined with organic manure (T20, T40, T60, T80) and sole organic manure (OM) reduced plant growth performance and yield. Growth traits and yield exhibited a gradual declining trend as the proportion of chemical-fertilizer substitution by organic manure increased. Relative to the CF treatment, stem length, internode length and number of fresh stems decreased significantly by 9.7%-27.9%, 14.9%-30.7% and 18.9%-42.4% under T40, T60, T80 and OM treatments; stem diameter, leaf number, chlorophyll content, yield and dry-matter conversion rate also declined significantly by 11.5%-18.5%, 7.5%-24.1%, 7.8%-22.1%, 4.2%-32.1% and 3.2%-5.6%, respectively (*P*<0.05).

### 3.2. Effects of Different Fertilization Treatments on Quality of *D. officinale*

Different fertilization treatments significantly affected the quality of *D. officinale*. All fertilized treatments exhibited higher contents of dendrobine, polysaccharides, polyphenols, flavonoids, alcohol-soluble extracts and protein compared with the non-fertilization control (Figure 1). Sole chemical fertilizer application, reduced chemical fertilizer combined with organic manure, and sole organic fertilizer application all significantly altered the contents of dendrobine, polyphenols and alcohol-soluble extracts. Relative to the non-fertilization treatment (CK), these components increased significantly by 21.2%-83.7%, 14.0%-49.1% and 10.8%-26.9%, respectively (*P*<0.05) (Figure 1a, b, c).

**Figure 1.**
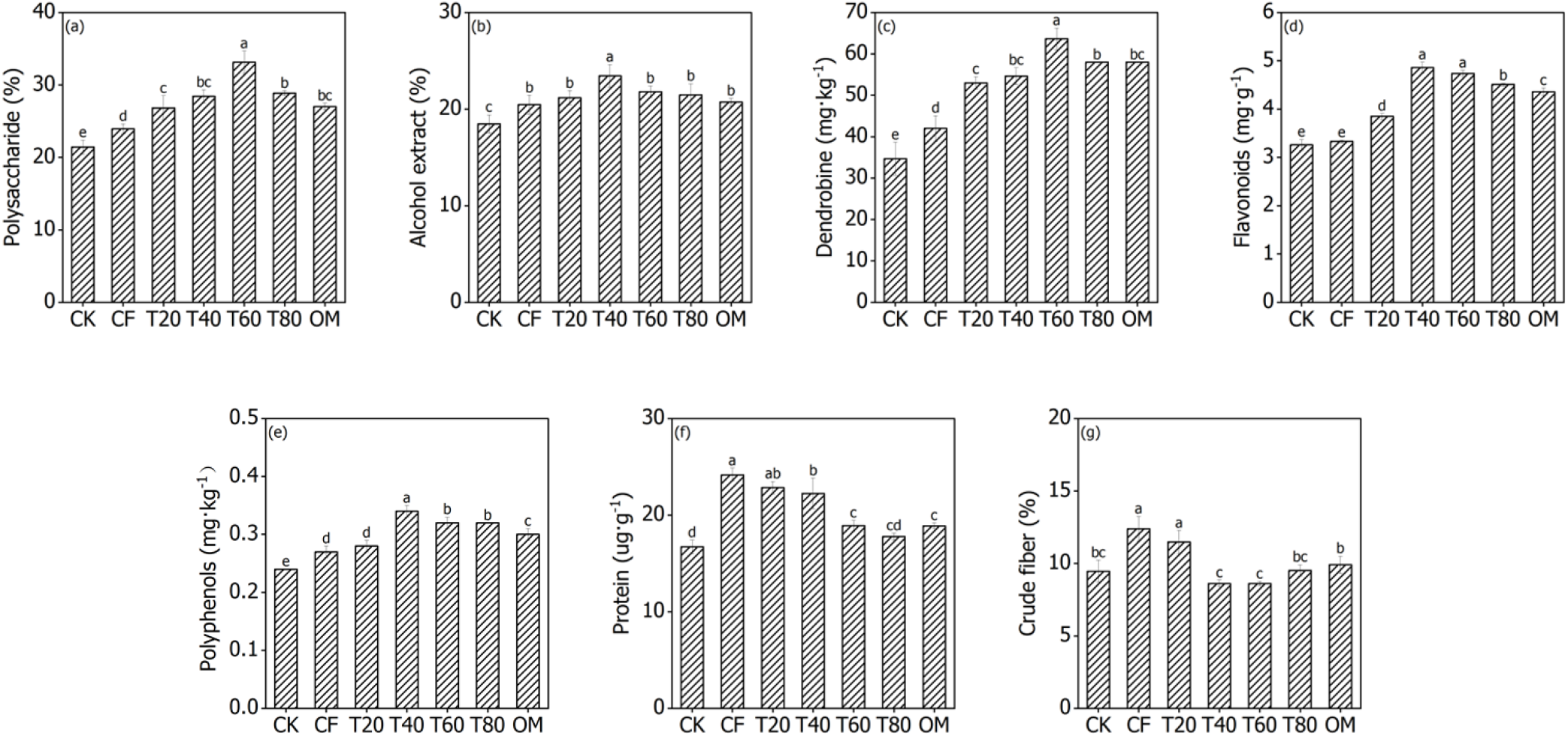
Effects of different fertilization treatments on quality of *D. officinale*. CK: No fertilization; CF: Sole chemical fertilizer; T20: 20% organic fertilizer + 80% chemical fertilizer; T40: 40% organic fertilizer + 60% chemical fertilizer; T60: 60% organic fertilizer + 40% chemical fertilizer; T80: 80% organic fertilizer + 20% chemical fertilizer; OM: Sole organic fertilizer. Values are means ± standard deviation (n = 3). Values with the same lower-case letters are not significantly different among different treatments at the 5% level by the LSD. The same below.

In addition, compared with CK, polysaccharide contents in T20, T40, T60, T80 and OM treatments increased significantly by 25.1%, 32.6%, 54.5%, 34.6% and 26.1% (*P*<0.05); flavonoid contents increased significantly by 18.1%, 49.1%, 45.6%, 38.6% and 33.8% (*P*<0.05). Protein contents in CF, T20, T40, T60 and OM treatments rose significantly by 44.6%, 36.6%, 33.1%, 13.1% and 12.9%. Crude-fiber contents in CF and T20 treatments increased significantly by 31.0% and 21.3% (*P*<0.05) (Figure 1).

Quality indicators of *D. officinale* were significantly modified by combined application of reduced chemical fertilizer and organic manure (T20, T40, T60, T80) as well as sole organic manure treatment (OM). As the proportion of chemical-fertilizer reduction increased, the quality of *D. officinale* showed a trend of first increasing and then decreasing. Compared with the CF treatment, dendrobine, polysaccharide and flavonoid contents in T20, T40, T60, T80 and OM treatments increased significantly by 26.2%-51.6%, 12.1%-38.4% and 15.5%-45.8%, respectively. Polyphenol contents in T40, T60, T80 and OM treatments increased significantly by 24.3%, 16.8%, 16.4% and 8.3%, whereas protein contents decreased significantly by 7.9%, 21.8%, 26.5% and 21.9%, and crude-fiber contents decreased significantly by 30.4%, 30.6%, 23.2% and 20.0%. Notably, alcohol-soluble extracts in the T40 treatment increased significantly by 14.5% (*P*<0.05).

### 3.3. Effects of Different Fertilization Treatments on Amino-Acid Composition of *D. officinale*

Amino acids are fundamental substances for all living organisms. In plants, amino acids are synthesized via biochemical reactions driven by various enzymes and enzyme systems during photosynthesis. For plants themselves, amino acids can promote root development, seedling vigour, enhance leaf photosynthetic capacity and resistance to pests and diseases, thereby improving yield. For human beings, plant-derived amino acids supply essential amino acids that cannot be synthesized de novo by the human body and play an indispensable role in life activities. The determination results of amino-acid composition of *D. officinale* under different fertilization treatments are shown in Table 2. As presented in Table 2, all fertilization treatments yielded complete amino-acid profiles, with a total of 17 amino acids detected in all samples.

**Table 2.** Effects of Different Fertilization Treatments on Amino-Acid Composition of *D. officinale*.

| Amino acid |  | CK | CF | T20 | T40 | T60 | T80 | OM |
|---|---|---|---|---|---|---|---|---|
| Essential amino acid (%) | Threonine | 0.107±0.01d | 0.18±0.006c | 0.199±0.023c | 0.26±0.016b | 0.308±0.013a | 0.12±0.01d | 0.099±0.02d |
|  | Phenylalanine | 0.137±0.015e | 0.222±0.008c | 0.238±0.015c | 0.308±0.027b | 0.341±0.014a | 0.168±0.01d | 0.092±0.011f |
|  | Lysine | 0.180±0.006c | 0.345±0.015b | 0.349±0.055b | 0.418±0.015a | 0.437±0.025a | 0.32±0.024b | 0.186±0.018c |
|  | Valine | 0.093±0.005f | 0.194±0.004d | 0.236±0.016c | 0.276±0.015b | 0.314±0.018a | 0.154±0.017e | 0.092±0.017f |
|  | Methionine | 0.015±0.004e | 0.044±0.007c | 0.049±0.002bc | 0.056±0.002a | 0.053±0.004ab | 0.025±0.002d | 0.016±0.003e |
|  | Isoleucine | 0.103±0.008e | 0.152±0.009d | 0.258±0.007c | 0.299±0.018b | 0.322±0.007a | 0.144±0.008d | 0.097±0.008e |
|  | Leucine | 0.165±0.017e | 0.418±0.01b | 0.588±0.035a | 0.622±0.02a | 0.607±0.053a | 0.342±0.022c | 0.268±0.022d |
| Nonessential amino acid (%) | Aspartic acid | 0.167±0.016d | 0.334±0.022c | 0.584±0.046b | 1.003±0.026a | 1.08±0.073a | 0.641±0.081b | 0.411±0.028c |
|  | Tyrosine | 0.06±0.01e | 0.159±0.01c | 0.165±0.013bc | 0.192±0.012b | 0.22±0.02a | 0.119±0.028d | 0.07±0.004e |
|  | Serine | 0.133±0.01bc | 0.137±0.005bc | 0.149±0.011b | 0.243±0.008a | 0.256±0.004a | 0.121±0.012cd | 0.108±0.017d |
|  | Glutamic acid | 0.176±0.011f | 0.506±0.028d | 0.645±0.08c | 0.983±0.071a | 0.863±0.093b | 0.425±0.095de | 0.355±0.05e |
|  | Glycine | 0.145±0.012e | 0.023±0.001f | 0.175±0.017d | 0.282±0.006b | 0.369±0.01a | 0.227±0.015c | 0.157±0.009de |
|  | Alanine | 0.142±0.01e | 0.188±0.01d | 0.229±0.025c | 0.301±0.019a | 0.267±0.012b | 0.162±0.02de | 0.152±0.009e |
|  | Cystine | 0.043±0.001ab | 0.031±0.001d | 0.034±0.004bc | 0.042±0.002ab | 0.045±0.002a | 0.034±0.012bc | 0.031±0.004d |
|  | Histidine | 0.031±0.003f | 0.076±0.007d | 0.161±0.01b | 0.181±0.002a | 0.086±0.007c | 0.044±0.002e | 0.023±0.003f |
|  | Arginine | 0.206±0.018e | 0.296±0.007c | 0.373±0.01b | 0.437±0.005a | 0.424±0.023a | 0.259±0.008d | 0.24±0.017d |
|  | Proline | 0.108±0.013f | 0.249±0.016c | 0.273±0.01b | 0.315±0.008a | 0.32±0.018a | 0.176±0.011d | 0.138±0.007e |
| Essential amino acid, EAA (%) |  | 0.801±0.031f | 1.555±0.011d | 1.916±0.02c | 2.239±0.085b | 2.383±0.012a | 1.273±0.061e | 0.85±0.064f |
| Nonessential amino acid, NEAA (%) |  | 1.211±0.018f | 1.998±0.03d | 2.79±0.086b | 3.978±0.056a | 3.931±0.06a | 2.207±0.004c | 1.685±0.064e |
| Total amino acid, TAA (%) |  | 2.012±0.048e | 3.553±0.038c | 4.706±0.105b | 6.217±0.03a | 6.314±0.048a | 3.48±0.057c | 2.534±0.113d |
| Eaa/TAA |  | 0.398±0.006b | 0.438±0.003a | 0.407±0.005b | 0.36±0.012d | 0.377±0.005c | 0.366±0.011cd | 0.335±0.014f |
| Neaa/TAA |  | 0.602±0.006d | 0.562±0.003e | 0.593±0.005d | 0.64±0.012b | 0.623±0.005c | 0.634±0.011bc | 0.665±0.014a |

Different fertilization treatments significantly affected the total amino-acid content and individual amino-acid concentrations of *D. officinale* (Table 2). In terms of total content and proportion of amino acids, sole chemical fertilizer application, reduced chemical fertilizer combined with organic manure, and sole organic fertilizer application significantly altered total amino-acid content and total non-essential amino-acid content. Compared with the non-fertilization control (CK), these two parameters increased significantly by 25.9%-213.8% and 39.1%-228.5%, respectively (*P*<0.05). Except for the OM treatment, essential amino-acid content increased significantly by 58.8%-197.4%. Regarding the proportional relationships among essential amino acids, non-essential amino acids and total amino acids, the ratios of EAA/TAA, NEAA/TAA and EAA/NEAA ranged from 0.355 to 0.438, 0.562 to 0.665, and 0.504 to 0.778 across fertilization treatments, respectively. Compared with CK, the CF treatment significantly raised EAA/TAA and EAA/NEAA by 9.9% and 17.7%, while significantly decreasing NEAA/TAA by 6.6%. For T40, T60, T80 and OM treatments, EAA/TAA and EAA/NEAA declined significantly by 5.2%-15.8% and 8.3%-23.8%, whereas NEAA/TAA increased significantly by 3.4%-10.5% (*P*<0.05).

For amino-acid composition, fertilization generally increased the concentrations of most individual amino acids, except for serine and cystine. As chemical-fertilizer dosage was reduced and organic-manure proportion increased, the content of each amino-acid component exhibited a trend of first increasing and then decreasing. Relative to CK, the contents of leucine, aspartic acid, glutamic acid, arginine and proline under all fertilization treatments increased significantly by 62.9%-277.9%, 100.6%-547.8%, 101.8%-458.6%, 16.6%-111.9% and 27.5%-196.4%, respectively. Furthermore, compared with CK, threonine and alanine contents increased significantly by 67.3%-187.1% and 32.5%-111.9% in CF, T20, T40 and T60 treatments. Phenylalanine, lysine, valine, methionine, isoleucine, tyrosine and histidine increased significantly by 22.6%-148.6%, 77.9%-142.8%, 64.4%-236.3%, 65.7%-270.0%, 38.9%-211.8%, 97.8%-266.2% and 40.0%-481.6% in CF, T20, T40, T60 and T80 treatments, respectively (*P*<0.05). In addition, compared with CK, phenylalanine content decreased significantly by 33.1% under OM. Serine content increased significantly by 82.7% and 92.2% in T40 and T60, but decreased significantly by 9.0% and 18.9% in T80 and OM. Glycine content dropped significantly by 84.5% under CF, whereas it increased significantly by 20.7%, 94.4%, 154.2% and 56.4% in T20, T40, T60 and T80. Cystine content decreased significantly by 28.1% under CF and increased significantly by 26.6% under OM (*P*<0.05). No significant differences were observed for other treatments.

Combined application of reduced chemical fertilizer with organic manure (T20, T40, T60, T80) and sole organic manure (OM) mostly exerted significant influences on total amino-acid content and individual amino-acid concentrations of *D. officinale*. With increasing proportion of chemical-fertilizer substitution, total amino-acid content and each amino-acid component generally showed an upward-then-downward trend, with optimal performance observed in T40 and T60 treatments. In respect of total content and ratios of amino acids, compared with CF, essential amino-acid content increased significantly by 23.2%, 44.0% and 53.3% in T20, T40 and T60, while decreasing significantly by 18.2% and 45.4% in T80 and OM. Non-essential amino-acid content increased significantly by 39.6%, 99.1%, 96.7% and 10.5% in T20, T40, T60 and T80, and decreased significantly by 15.7% under OM. Total amino-acid content increased significantly by 32.5%, 75.0% and 77.7% in T20, T40 and T60, and decreased significantly by 28.7% under OM. Moreover, compared with CF, treatments of reduced chemical fertilizer combined with organic manure (T20, T40, T60, T80) and sole organic manure (OM) significantly reduced EAA/TAA and EAA/NEAA by 6.9%-23.4% and 11.7%-35.2%, and significantly elevated NEAA/TAA by 5.4%-18.2% (*P*<0.05).

In terms of amino-acid composition, T20, T40 and T60 increased the contents of all 17 amino acids. By contrast, T80 and OM reduced 14 amino-acid components except aspartic acid, glycine and cystine. Compared with CF, the T20 treatment significantly increased valine, isoleucine, leucine, aspartic acid, glutamic acid, glycine, alanine, cystine, histidine, arginine and proline by 21.7%, 69.9%, 40.8%, 74.8%, 27.5%, 677.1%, 21.6%, 11.3%, 113.5%, 26.2% and 9.7%, respectively. T40 and T60 significantly enhanced all amino-acid components, with increments ranging from 20.6% to 1151.7% and from 20.1% to 1536.9%, respectively. Under T80, threonine, phenylalanine, valine, methionine, leucine, aspartic acid, glycine, cystine, histidine, arginine and proline increased significantly by 33.4%, 24.3%, 20.7%, 43.3%, 18.1%, 18.2%, 25.4%, 907.5%, 11.9%, 42.4%, 12.6%, 29.5% and 10.5%, while tyrosine decreased significantly by 91.7%. For the OM treatment, threonine, phenylalanine, lysine, valine, methionine, isoleucine, leucine, tyrosine, serine, glutamic acid, alanine, histidine, arginine and proline decreased significantly by 44.8%, 58.7%, 46.2%, 52.3%, 64.5%, 36.3%, 35.8%, 56.3%, 21.1%, 29.8%, 19.1%, 69.6%, 18.7% and 44.8%, respectively; glycine content increased significantly by 594.5%. No significant differences were detected among other comparisons.

It can be concluded that although reduced chemical-fertilizer combined with organic-manure application lowered the yield of *D. officinale*, amino-acid contents were markedly improved under appropriate substitution ratios (T20, T40 and T60). From the perspective of amino-acid profiles, the rank order of amino-acid types and concentrations was largely consistent across fertilization treatments. Aspartic acid possessed the highest content, followed by glutamic acid, leucine, arginine and lysine. Aspartic acid and glutamic acid are characteristic umami-taste amino acids, jointly accounting for more than 25% of total amino-acids and contributing to flavour regulation.

Among the 17 amino acids detected in *D. officinale* under all fertilization treatments, four umami-taste amino acids DAA (Asp, Glu, Gly, Ala), eight bitter-taste amino acids BAA (Val, Met, Ile, Leu, Tyr, Phe, His, Arg), and nine medicinal amino acids MAA (Asp, Glu, Gly, Arg, Met, Phe, Leu, Lys, Tyr) were identified. The contents of bitter-taste amino acids, umami-taste amino acids and medicinal amino acids ranged from 63.3% to 258.0%, 81.1% to 237.1%, and 129.9% to 448.2%, respectively. Different fertilization treatments significantly affected the contents of these three categories of amino acids. Compared with the non-fertilization control (CK), the contents of bitter-taste amino acids, umami-taste amino acids and medicinal amino acids in fertilized *D. officinale* increased significantly by 66.8%-309.5%, 10.7%-192.3% and 40.4%-245.2%, respectively. Relative to the CF treatment, bitter-taste amino acids increased significantly by 55.4%, 144.5%, 145.5% and 38.5% under T20, T40, T60 and T80; umami-taste amino acids increased by 32.6%, 52.0% and 51.7% under T20, T40 and T60, but decreased significantly by 19.7% and 42.5% under T80 and OM; medicinal amino acids increased significantly by 35.2%, 84.5%, 89.3% and 6.8% under T20, T40, T60 and T80, and decreased significantly by 23.0% under OM (Figure 2). Notably, medicinal amino acids exhibited relatively high content and proportion. The ratio of medicinal amino acids to total amino acids ranged from 64.5% to 72.6% across treatments, with an average value of 69.3%. The ranking of medicinal-amino-acid content from high to low was as follows: T60 = T40 > T20 > T80 > CF > OM > CK (Figure 2). These results indicated that *Dendrobium* plants are rich in medicinal amino acids, and rational combined application of reduced chemical fertilizer with organic manure can further improve the accumulation of medicinal amino-acid components.

**Figure 2.**
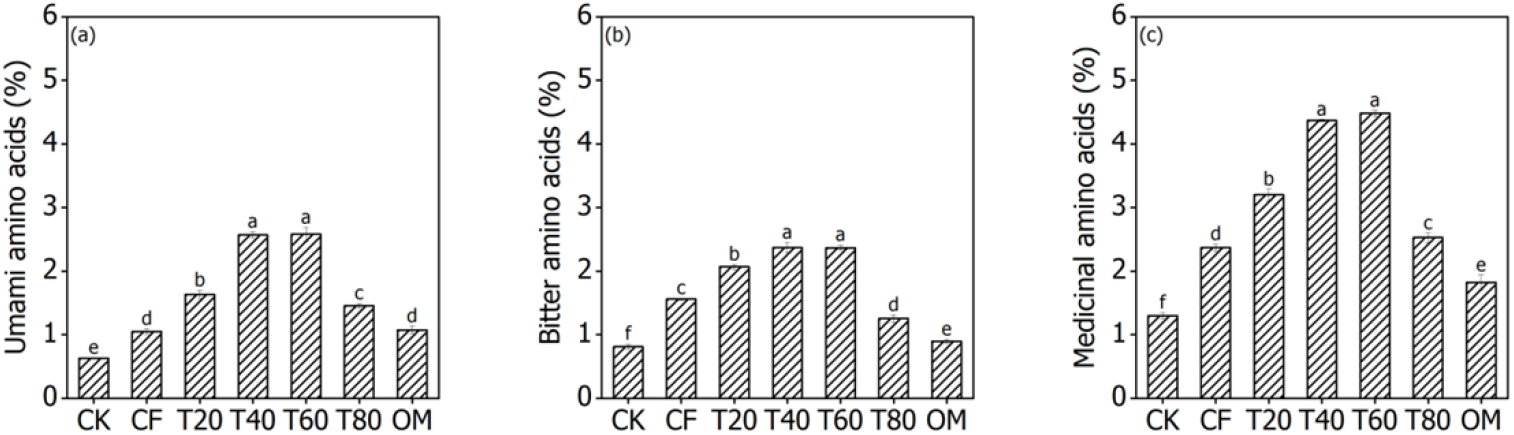
Contents of different categories of amino acids under various fertilization treatments

### 3.4. Correlation Analysis of Various Indicators of *D. officinale*

Correlation analysis was performed on multiple indicators of *D. officinale* (Figure 3), and significant correlations were observed among these parameters. Pearson correlation analysis revealed that growth-related traits including stem length, stem diameter, internode length, leaf number, chlorophyll content, number of fresh stems, yield and dry-matter conversion rate showed highly significant positive correlations with one another. Except for protein and crude fiber, quality indicators such as dendrobine, polysaccharides, polyphenols, flavonoids and alcohol-soluble extracts also exhibited highly significant positive correlations. Highly significant positive correlations were also detected among essential amino acids, non-essential amino acids, total amino acids, umami-taste amino acids, bitter-taste amino acids and medicinal amino acids. Furthermore, quality indicators (dendrobine, polysaccharides, polyphenols, flavonoids, alcohol-soluble extracts) were significantly or highly significantly positively correlated with essential amino acids, non-essential amino acids, total amino acids, umami-taste amino acids, bitter-taste amino acids and medicinal amino acids. In addition, protein showed highly significant positive correlations with all growth indicators of *D. officinale*. Crude fiber was significantly positively correlated with most growth indicators but negatively correlated with quality indicators. Leaf number and yield were both highly significantly positively correlated with all categories of amino acids (Figure 3).

**Figure 3.**
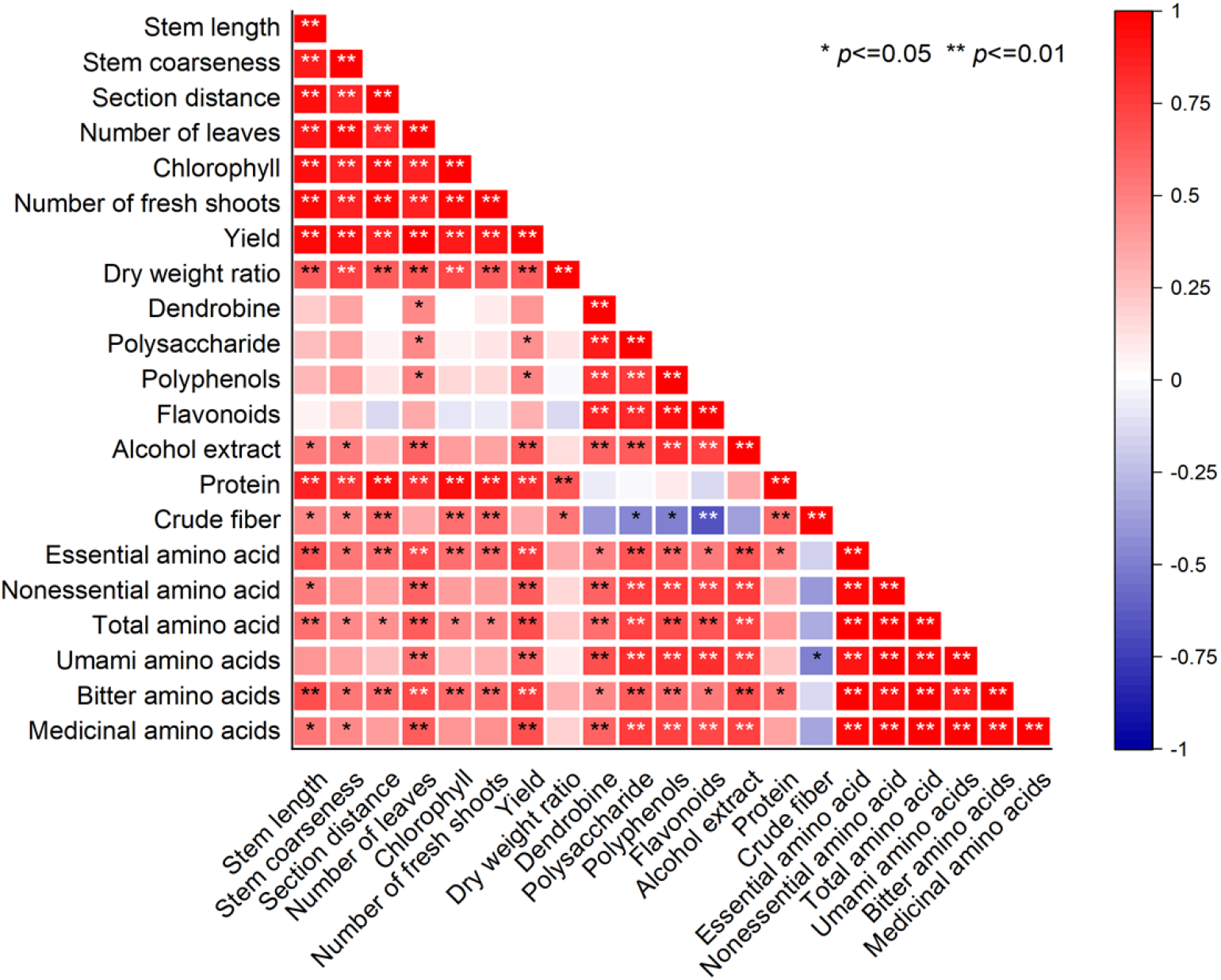
Correlation coefficients among various indicators of *D. officinale*

### 3.5. Comprehensive Evaluation of the Effects of Different Fertilization Treatments on Growth Performance and Quality of *D. officinale*

The membership-function method was adopted to comprehensively evaluate the effects of various fertilization treatments on the growth performance and quality of *D. officinale* (Table 3). The results showed that the comprehensive evaluation values under different fertilization treatments were ranked in the order: T40 > T60 > CF > T20 > T80 > OM > CK. The comprehensive evaluation values of sole chemical fertilizer application, combined application of reduced chemical fertilizer with organic manure, and sole organic manure application were all higher than that of the non-fertilization control.

**Table 3.** Membership-function-based comprehensive evaluation of growth, yield and nutraceutical quality of *D. officinale* under different fertilization treatments.

| Index | CK | CF | T20 | T40 | T60 | T80 | OM | Vj | Wj |
|---|---|---|---|---|---|---|---|---|---|
| Stem length | 0.004 | 0.051 | 0.047 | 0.039 | 0.032 | 0.028 | 0.016 | 0.514 | 0.055 |
| Stem diameter | 0.002 | 0.047 | 0.034 | 0.032 | 0.031 | 0.029 | 0.026 | 0.451 | 0.048 |
| Internode length | 0.003 | 0.061 | 0.052 | 0.038 | 0.028 | 0.018 | 0.016 | 0.653 | 0.070 |
| Number of leaves | 0.001 | 0.047 | 0.044 | 0.042 | 0.038 | 0.030 | 0.028 | 0.456 | 0.049 |
| Chlorophyll | 0.004 | 0.066 | 0.048 | 0.044 | 0.029 | 0.023 | 0.014 | 0.633 | 0.067 |
| Number of fresh shoots | 0.003 | 0.058 | 0.051 | 0.038 | 0.028 | 0.027 | 0.013 | 0.600 | 0.064 |
| Dry weight ratio | 0.009 | 0.053 | 0.030 | 0.024 | 0.032 | 0.016 | 0.027 | 0.552 | 0.059 |
| Yield | 0.001 | 0.049 | 0.045 | 0.044 | 0.039 | 0.032 | 0.023 | 0.475 | 0.051 |
| Dendrobine | 0.004 | 0.016 | 0.033 | 0.036 | 0.050 | 0.041 | 0.041 | 0.493 | 0.053 |
| Polysaccharide | 0.004 | 0.015 | 0.027 | 0.034 | 0.054 | 0.036 | 0.028 | 0.554 | 0.059 |
| Polyphenols | 0.001 | 0.021 | 0.022 | 0.059 | 0.047 | 0.047 | 0.034 | 0.589 | 0.063 |
| Flavonoids | 0.004 | 0.007 | 0.026 | 0.064 | 0.060 | 0.051 | 0.045 | 0.633 | 0.067 |
| Alcohol extract | 0.005 | 0.020 | 0.025 | 0.042 | 0.030 | 0.027 | 0.022 | 0.480 | 0.051 |
| Protein | 0.025 | 0.062 | 0.051 | 0.047 | 0.021 | 0.012 | 0.021 | 0.630 | 0.067 |
| Crude fiber | 0.037 | 0.008 | 0.017 | 0.045 | 0.045 | 0.036 | 0.032 | 0.445 | 0.047 |
| Total amino acids | 0.001 | 0.029 | 0.050 | 0.077 | 0.079 | 0.027 | 0.010 | 0.750 | 0.080 |
| Comprehensive evaluation value | 0.111 | 0.657 | 0.648 | 0.748 | 0.682 | 0.514 | 0.420 |  |  |
| Ranking | 7 | 3 | 4 | 1 | 2 | 5 | 6 |  |  |

Among all treatments, the T40 treatment (40% organic manure substitution for chemical fertilizer) obtained the highest comprehensive evaluation value of 0.748 for growth performance and quality, followed by the T60 treatment (60% organic manure substitution for chemical fertilizer) with a value of 0.682. The sole organic-manure treatment (OM) had the lowest comprehensive evaluation value of 0.420.

These findings indicated that fertilization generally improved the comprehensive performance of *D. officinale* in terms of growth and quality. Chemical fertilizer mainly promoted plant growth traits and yield, whereas organic manure predominantly improved quality attributes. Reasonable allocation of chemical and organic fertilizer dosages could achieve an optimal balance between yield and quality.

## 4. Discussion

### 4.1 Effects of Different Fertilization Treatments on Growth Traits and Yield of D. officinale

The present study demonstrated that fertilization could promote the growth and improve quality of *D. officinale*, achieving high yield and good quality simultaneously[35-36]. In this experiment, fertilization significantly improved growth indices including stem length, stem diameter, internode length, leaf number, chlorophyll content and number of fresh stems, and increased yield, which was consistent with the findings reported by Wu et al. Their research confirmed that stem length, stem diameter, fresh weight per clump, water content, crude-fiber content, alkaloid content, polysaccharide content, total flavonoid content, alcohol-soluble extract content and protein content of fertilized *D. officinale* were all significantly higher than those under non-fertilization treatment.

In addition, with the increasing substitution ratio of organic manure for chemical fertilizer, growth traits and yield of *D. officinale* gradually declined. This result agreed with the conclusion of Liu et al.[37]. Chemical fertilizer exhibited prominent advantages in promoting vegetative growth and biomass production of *D. officinale*, while organic manure showed weaker promoting effects on plant vegetative growth. Chemical fertilizer can rapidly release mineral nutrients such as nitrogen, phosphorus and potassium to satisfy the nutritional demands for rapid tillering and stem elongation of *D. officinale*. It raises leaf chlorophyll concentration, enhances photosynthetic efficiency, and consequently increases yield, which is in accordance with previous studies[24]. Zhang et al. applied special controlled-release fertilizer for orchids to *Dendrobium denneanum*, and observed significant promotion in stem growth rather than leaf growth; stem length, stem diameter and internode number showed consistent increasing trends[38].

Under the experimental conditions of this study, the yield of T20 and T40 treatments only slightly decreased by 4.2%-6.0% compared with CF and remained at a relatively high level. Nevertheless, when the organic-manure substitution ratio reached 80% (T80) or under sole organic-manure treatment (OM), yield decreased significantly by 20.8% and 32.1% relative to CF, respectively. This experiment adopted an equal-nitrogen substitution design. Nutrients from organic manure are released slowly. Even with identical total nitrogen input, high-proportion organic-manure treatments provide insufficient available mineral nitrogen during the vigorous-growth stage, failing to meet the demand for rapid vegetative growth of *D. officinale*. This further inhibits tillering, stem elongation and fresh-stem yield formation. These observations are similar to those reported by Kong et al.[39] in fertilizer trials on *D. officinale*. Although high-rate organic manure can improve substrate properties, insufficient supply of readily available nutrients restricts above-ground biomass accumulation[40].

All combined fertilization treatments in this study produced significantly higher yield than the non-fertilization control, indicating that reduced chemical fertilizer combined with organic manure was still superior to non-fertilized cultivation. In medicinal plant cultivation, sole chemical fertilizer application pursuing high yield usually leads to poor quality, whereas sole organic-manure application causes yield loss. This represents a practical contradiction in commercial production of *D. officinale*. Therefore, an appropriate organic-manure substitution ratio should be determined within an acceptable yield-loss range to realize coordinated improvement of yield and quality. The yield of T40 treatment in this trial was only 5.7% lower than that of CF, showing a minor yield reduction and providing a feasible option for balancing yield performance. It should be noted that this experiment lasted for one annual growth cycle. Substrate physicochemical properties will continuously change along with long-term decomposition of organic manure. Further long-term field trials are required to clarify the persistent effects of different fertilizer combinations on growth and yield of *D. officinale* under multi-year continuous cultivation.

### 4.2 Effects of Different Fertilization Treatments on Quality of D. officinale

The medicinal value of *D. officinale* depends on the contents of bioactive components including dendrobine, polysaccharides, polyphenols, flavonoids and alcohol-soluble extracts, which serve as core indices for evaluating medicinal material quality[2-4]. The present study showed that the contents of dendrobine, polysaccharides, polyphenols, flavonoids, alcohol-soluble extracts and protein under all fertilization treatments were higher than those under CK. With the increase in the organic-manure substitution ratio for chemical fertilizer, the medicinal quality indices of *D. officinale* generally exhibited a trend of first increasing and then decreasing. The T40 and T60 treatments produced the most prominent improvement in polysaccharide, dendrobine, flavonoid and polyphenol contents. Compared with sole chemical-fertilizer treatment (CF), T40 significantly increased dendrobine, polysaccharides and flavonoids by 47.4%, 32.8% and 45.8%, respectively. These findings support previous reports that organic manure facilitates the accumulation of secondary-metabolite bioactive substances in *D. officinale* and can markedly elevate the contents of polysaccharides, alkaloids and flavonoids relative to sole chemical fertilizer application[24,40].

In addition to macronutrients such as nitrogen, phosphorus and potassium, organic manure contains humic acids, small-molecule organic acids, trace elements and microbial metabolites. On the one hand, it ameliorates the micro-environment of the cultivation substrate; on the other hand, it induces the expression of secondary-metabolic pathways in *D. officinale*, promoting the synthesis and accumulation of defensive and medicinal bioactive substances such as polyphenols, flavonoids and alkaloids[41]. Nevertheless, quality indices declined under excessive organic-manure substitution (T80, OM). It is speculated that excessive organic nutrient supply disturbs the carbon-nitrogen metabolic balance of plants, and more carbon skeletons are consumed by substrate microorganisms, thereby inhibiting the biosynthesis of secondary metabolites. The polysaccharide and alcohol-soluble extract contents in all treatments of this study complied with the quality criteria for *D. officinale* specified in the *Pharmacopoeia of the People’s Republic of China* (2020 Edition). Combined fertilization treatments exhibited obvious advantages in quality improvement. Although CF achieved the highest yield, its contents of medicinal bioactive components including polysaccharides and dendrobine remained at relatively low levels, indicating that cultivation relying solely on chemical fertilizer carries risks of insufficient medicinal quality for *D. officinale*.

In the present study, the variation trends of protein and crude fiber were not fully consistent with those of polysaccharides and dendrobine. Protein content decreased with the reduction in chemical-fertilizer proportion, with the maximum value observed in CF. Higher crude-fiber contents were also recorded in CF and T20, whereas crude fiber decreased under treatments with high organic-manure proportion. Correlation analysis revealed that protein was highly significantly positively correlated with growth indices but negatively correlated with most bioactive quality indices. This suggests that protein and crude fiber are more closely associated with the vigour of vegetative growth, whereas dendrobine, polysaccharides and flavonoids represent medicinal secondary quality. These two groups of parameters exhibit distinct response patterns to fertilizer nutrients. This result is similar to the conclusion reported by Wang et al.[42] regarding the differentiated responses of different nutritional components to nutrient conditions in mycorrhizal-inoculated *D. officinale*. In research on fertilization techniques for *D. officinale*, quality assessment should not rely on a single indicator; instead, yield and multiple bioactive components should be comprehensively taken into consideration.

### 4.3 Effects of Different Fertilization Treatments on Amino-Acid Composition of D. officinale

Amino acids are important nutritional components of *D. officinale*. Among them, umami-taste, bitter-taste and medicinal amino acids are closely associated with its dietary and health-promoting values[8-9]. All 17 amino acids were detected across the seven treatments in this experiment. Fertilization significantly increased the contents of total amino acids, essential amino acids, medicinal amino acids, umami-taste amino acids and bitter-taste amino acids. With the increase of organic-manure substitution ratio, total amino acids, essential amino acids, medicinal amino acids and umami-taste amino acids exhibited a trend of first increasing and then decreasing. The T40 and T60 treatments achieved the best comprehensive performance in amino-acid profiles. Compared with CF, T40 increased total amino acids by 75.0 % and medicinal amino acids by 84.5 %. Nevertheless, further elevation of organic-manure substitution ratio led to a remarkable decline in various amino-acid contents under T80 and OM; the total amino-acid content under OM was even lower than that under CF.

Organic manure contains abundant organic-nitrogen substrates such as amino acids and polypeptides, which can be directly absorbed and utilized by the root system of orchid plants. Meanwhile, functional microorganisms including *Bacillus subtilis* in the substrate can accelerate the mineralization of organic matters, release small-molecular amino-acid substances, facilitate the synthesis and accumulation of amino acids in plants, and consequently raise the contents of essential and medicinal amino acids. The organic manure adopted in this study was fermented from mugwort straw, earthworm cast and *Pleurotus eryngii* residue, compounded with extracts of *D. officinale* and mugwort, and inoculated with *Bacillus subtilis*. It can supply abundant organic nitrogen sources for *D. officinale*, which forms the material basis for the substantial improvement of amino-acid components in T40 and T60 treatments. However, under high-substitution treatment T80 and sole organic-manure treatment OM, the supply of readily-available mineral nitrogen was insufficient, restricting total nitrogen uptake by plants and resulting in shortage of precursors for amino-acid synthesis, thereby decreasing total amino-acid content.

Aspartic acid and glutamic acid occupied prominent proportions in the amino-acid profiles of *D. officinale* in the present study. As umami-taste amino acids, they jointly accounted for more than 25 % of total amino acids and determine the edible flavour of *D. officinale*. Medicinal amino-acid components accounted for 64.5 %-72.6 % of total amino acids, with an average value as high as 69.3 %, which constitutes the material foundation for the health-beneficial effects of *D. officinale*. T40 and T60 treatments exerted significant effects on enhancing medicinal and umami-taste amino-acids. It indicates that appropriate reduction of chemical fertilizer combined with organic manure can not only improve medicinal bioactive components, but also elevate the dietary nutritional value of *D. officinale*. Correlation analysis demonstrated that medicinal quality indices such as dendrobine, polysaccharides and flavonoids were significantly or highly significantly positively correlated with total amino acids and medicinal amino acids. This suggests synergistic regulation between the accumulation of secondary bioactive substances and amino-acid metabolism in *D. officinale*. This finding is consistent with the report by Wang et al.[43] on mycorrhizal-inoculated *D. officinale*, namely that cultivation conditions favourable for secondary metabolism can simultaneously promote amino-acid accumulation.

### 4.4 Insights from Comprehensive Evaluation on Yield-Quality Synergy of D. officinale

As a medicinal economic crop, *D. officinale* requires both acceptable fresh-stem yield and guaranteed intrinsic medicinal quality in practical production. Neither single yield indicator nor individual bioactive-component index can objectively judge the merits of fertilization regimes. The membership-function comprehensive evaluation method normalizes multi-dimensional parameters and enables integrated assessment covering growth, yield, quality and amino-acid profiles, which has been widely applied in screening cultivation measures for medicinal plants. In the present study, the ranking of comprehensive evaluation values was: T40 > T60 > CF > T20 > T80 > OM > CK.

The T40 treatment obtained the highest comprehensive evaluation value (0.748). Its yield decreased only slightly by 5.7% compared with CF which achieved the maximum yield, whereas dendrobine, polysaccharides, flavonoids, total amino acids and medicinal amino acids were all significantly enhanced. It realized maximum quality improvement with minimal yield loss. Although T60 also exhibited high quality indices, its yield decreased by 12.4% relative to CF, resulting in inferior comprehensive performance compared with T40. Sole chemical-fertilizer treatment (CF) ranked first in yield, yet its comprehensive evaluation value was lower than T40 and T60, owing to its insufficient medicinal bioactive components and total amino-acid content. Treatments with high organic-manure proportion (T80 and OM) showed partial improvement in quality indices but suffered considerable yield reduction, leading to unsatisfactory overall performance.

It can be concluded that sole chemical-fertilizer application focuses on yield promotion while organic manure prioritizes quality improvement, and a certain trade-off exists between them. Only an appropriate mixing ratio can achieve the balance of both targets. Under the experimental conditions of this study, 40% equal-nitrogen organic-manure substitution for chemical fertilizer (T40) was the optimal fertilization scheme for bed-cultivated *D. officinale* in greenhouse production to coordinate yield and quality. This experiment was carried out under specific greenhouse facilities, cultivation substrates and special organic manure in Ruili, Dehong. Multi-site verification is still required to confirm whether this ratio is fully applicable under alternative climatic conditions, substrate types and organic-manure raw-material sources. For practical popularization in production, further fine-tuning should be performed considering fertilizer cost, substrate fertility and cultivation duration.

## 5. Conclusions

In this study, a greenhouse bed-cultivation experiment was conducted to investigate the regulatory effects of reduced chemical fertilizer combined with organic manure on yield, quality and amino-acid composition of *D. officinale*. The results showed that fertilization significantly improved growth traits and enhanced yield and quality of *D. officinale*. Sole chemical-fertilizer treatment achieved the best vegetative growth and fresh-stem yield, yet the accumulation of medicinal bioactive components was limited. With the increase in organic-manure substitution ratio, stem length, stem diameter and yield of *D. officinale* exhibited a declining trend, whereas medicinal quality indices including dendrobine, polysaccharides and flavonoids followed a pattern of first increasing and then decreasing. Appropriate organic-manure substitution significantly increased the contents of total amino acids, essential amino acids, medicinal amino acids and umami-taste amino acids, with T40 and T60 showing the most pronounced improvement; high-rate organic-manure substitution led to a reduction in amino-acid contents. The average proportion of medicinal amino acids in total amino acids of *D. officinale* reached 69.3%. Most medicinal bioactive components were significantly or highly significantly positively correlated with amino-acid components, indicating a synergistic accumulation effect.

The 40% equal-nitrogen organic-manure substitution for chemical fertilizer (T40) obtained the highest comprehensive evaluation value. Under this treatment, yield only slightly decreased by 5.7% compared with sole chemical-fertilizer application, while the contents of dendrobine, polysaccharides, flavonoids and amino acids were significantly increased. Therefore, T40 was identified as the optimal fertilization mode for balancing yield and medicinal quality of *D. officinale* under the present experimental conditions.

## Author contributions

**Conceptualization:** Long Zhou, Hongbin Yang.

**Data curation:** Long Zhou.

**Formal analysis:** Long Zhou, Xudong Zhao.

**Investigation:** Long Zhou, Wenna Zhou, Hongbin Yang.

**Methodology:** Long Zhou.

**Validation:** Long Zhou, Huarong Liu.

**Writing – original draft:** Long Zhou.

**Writing – review & editing:** Long Zhou, Hongbin Yang.

## Data availability statement

All relevant data are within the paper and its Supporting Information files.

